# The *Arabidopsis* CYSTM α 5’UTR increases protein production from transgenes in plants and bacteria

**DOI:** 10.1101/2025.10.15.682649

**Authors:** Jasjyot Khanduja, Xingyu Wu, Jun Li, Iain Searle

## Abstract

Achieving high levels of transgene expression is often crucial for the effective production of high value pharmaceutical proteins and bioactive compounds. Here, we show a 79-nucleotide sequence of the 5’ UTR of the *Arabidopsis thaliana* cysteine-rich transmembrane module 1 *(CYSTM1*) significantly increases transient and stable transgene expression at the post-transcriptional level in multicellular plants, *A. thaliana* and *Nicotiana benthamiana*, in monocotyledon wheat germ extracts, and in the unicellular bacterium *Escherichia coli* by up to 7-fold. We also show the 79 nt *CYSTM* α sequence has 50 % higher transgene expression than the broadly used Tobacco Mosaic Virus omega (Ω) sequence in both *A. thaliana* and *N. benthamiana*. Taken together, this study provides a short transgene expression enhancer for multicellular and unicellular organisms.

## Introduction

Plant molecular farming (PMF) is a key area of plant biotechnology that focuses on the use of plants to produce beneficial products, including human therapeutic proteins (1-3). PMF has several advantages over conventional microbial or mammalian protein production systems, such as rapid scalability, cost-effectiveness, and increased biosafety (1-4). However, a major limitation of PMF is the relatively low and inconsistent expression of the transgenes. Improving transgene expression is therefore critical, as it not only increases product yield but also simplifies downstream purification, ultimately reducing overall production costs (5–10).

Transgene expression is regulated by different elements, including promoters, 5′ and 3′ untranslated regions (UTRs), introns, and terminators (5-8). These elements modulate transcriptional activity, mRNA stability, and translational efficiency (9, 10). In addition to these cis-regulatory elements, endogenous processes such as RNA silencing, histone modifications, and co-transcriptional RNA processing also play important roles in regulating transgene performance. It is well established that transgene components such as promoters, 5′ and 3′ untranslated regions (UTRs), introns, and terminators play crucial roles in regulating transcription efficiency, RNA stability, and translation rates (9-13). In addition, other cellular processes such as RNA silencing, histone modifications and co-transcriptional RNA modifications can also impact transgene expression (14-19). Notable examples of translational enhancers used in plant transgenes include the tobacco mosaic virus (TMV) omega (Ω) sequence, *AtADH* 5’UTR, synJ, *NtADH* 5’UTR and *OsADH* 5’ UTR (20-26).

Here we show a short, 79-nt sequence, derived from the 5′ UTR of the *A. thaliana* CYSTM1 gene (AT1G05340) that functions as a potent translational enhancer. This sequence significantly boosts transgene expression across multiple systems, including in planta (both in vivo and in vitro) and in bacteria (*E. coli*). Comparative analyses reveal that the CYSTM1 5′ UTR enhancer consistently outperforms the widely used TMV Ω enhancer, achieving up to a two-fold increase in transgene expression. These results suggest that the CYSTM1-derived element represents a promising alternative to traditional viral enhancers for improving gene expression in plant biotechnology and heterologous systems.

## Results

### Identification of a translation enhancer sequence in the 5’ UTR of *CYSTM* by transient expression in *N. benthamiana* leaves

Previously, our lab mapped RNA m^5^C sites on mRNA purified from *A. thaliana* seedlings (27). To test whether any of these modifications could impact translation, DNA fragments encoding the modified RNA regions were cloned into a *Firefly luciferase* (F-Luc) reporter either upstream (5′ UTR) or downstream (3′ UTR) of the F-Luc coding sequence (Supplementary Fig. 1). These constructs were agroinfiltrated into *N. benthamiana* leaves and assayed for bioluminescence. While most fragments showed bioluminescence similar to the control, one fragment, 5′ CYSTM1 α/ AT1G05340 was striking. 5′ CYSTM1 α increased bioluminescence when cloned in the 5′ UTR. We further investigated this enhanced bioluminescence as well as an established viral enhancer.

To compare the 79-nt 5′ CYSTM α enhancer with its native context and with a well-established viral enhancer, we cloned the full 110-nt CYSTM1 5′ UTR (pluc 5′ CYSTM1) and the 67-nt TMV Ω enhancer (pluc 5′ TMV Ω) upstream *Photinus pyralis* (firefly) luciferase gene and used the *Renilla reniformis* (sea pansy) luciferase as an internal control (Supplementary Fig 1). After *Agrobacterium*-mediated transient expression in *N. benthamiana* leaves, we observed a 4-fold higher firefly bioluminescence from both the TMV Ω enhancer (pluc 5’ TMV Ω) and *CYSTM1* 5’ UTR (pluc 5’ *CYSTM1*) compared to the no insert pluc control. (Fig. 1B).

**Figure 1:**
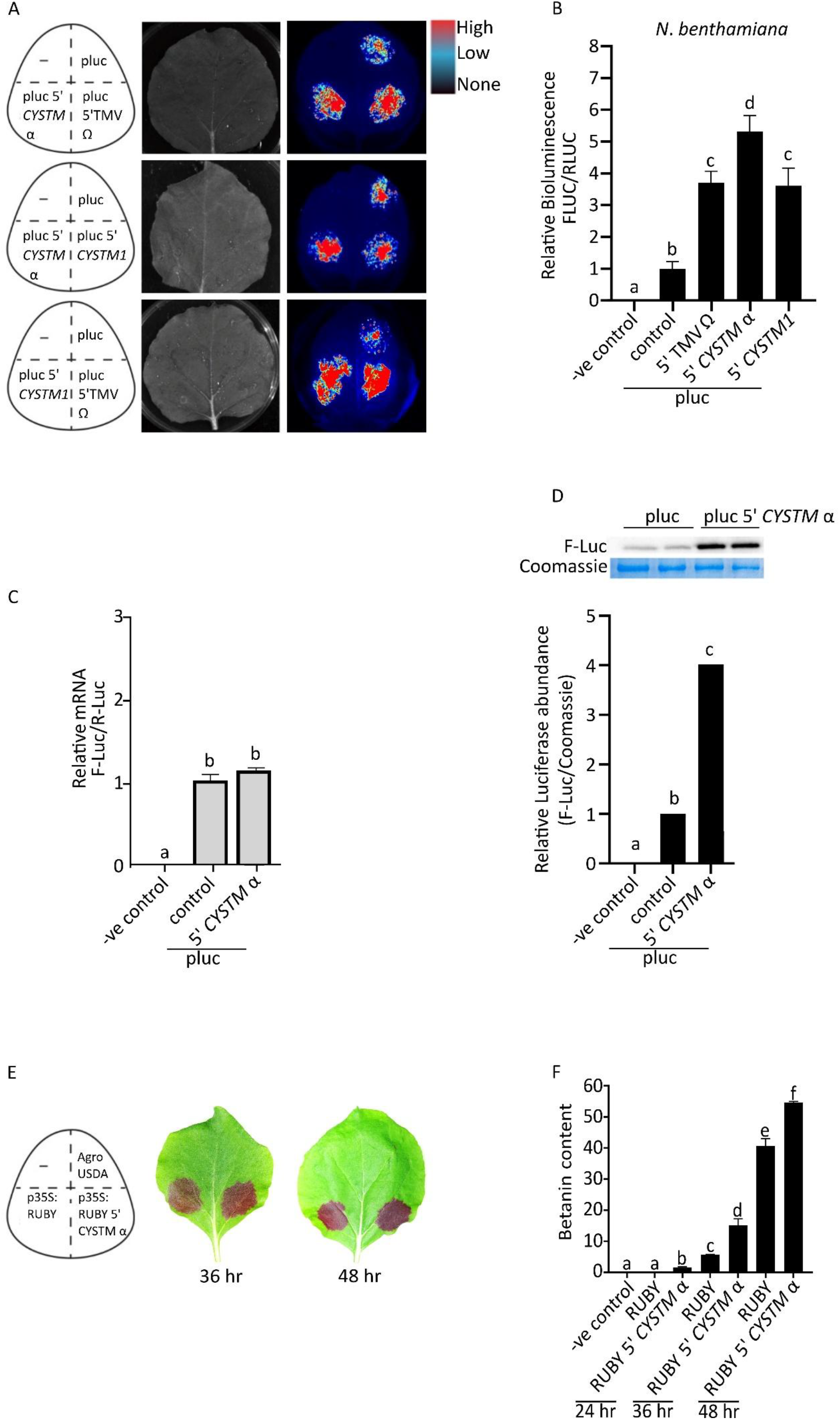
*CYSTM1* enhances dual luciferase and RUBY transgene expression in *N. benthamiana* leaves. **A)** Schematic representation of the treatments and bioluminescence imaging of *N. benthamiana* leaves 72 hours post-infiltration with either pluc (control), pluc 5’ *CYSTM* α (79 nt), pluc 5’ *CYSTM1* (110 nt), or pluc 5’ TMV Ω (67 nt) constructs. Imaging was conducted on a CCD camera (Biorad) using a chemi-high sensitivity. Left: Schematic of infiltration sites. Middle: Leaf under white light. Right: False-coloured bioluminescence images. Exposure time was adjusted per leaf as needed. **B)** Dual-Luciferase assay results for leaves expressing different constructs. Experiment conducted with 5 biological replicates and repeated thrice. Error bars indicate the standard error of the mean. Statistical significance denoted by letters (a-e) (P < 0.01, one-way ANOVA). **C)** qRT-PCR analysis of the *Firefly luciferase* mRNA levels. **D)** Semi-quantitative western blot analysis of LUCIFERASE. **E)** Temporal quantification of betanin in leaves expressing either the p35:RUBY control or 5’ *CYSTM* α constructs at 12, 24, and 36-hour time points. Statistical significance is indicated by letters (a-d) and was assessed using one-way ANOVA (P < 0.01).

In comparison, after agroinfiltrating the 79 nt *CYSTM1* 5’ UTR enhancer (pluc 5’ *CYSTM* α), we observed a 5.5-fold increase in bioluminescence compared to the control (Fig. 1 B). The short 79 nt sequence gave 30 % more firefly bioluminescence than the full-length *CYSTM1* 5’ UTR.

Next, we conducted qRT-PCR to quantify the firefly luciferase mRNA abundance (Fig 1 C), and we observed no difference in the mRNA abundances between the pluc control and the pluc 5’ *CYSTM* α (Fig. 1 C). Western blot and ImageJ analysis revealed approximately a four-fold higher LUCIFERASE protein from the pluc 5’ *CYSTM* α transgene compared to the control (Fig. 1 D). Together, these results suggest the pluc 5’ *CYSTM* α increases protein translation.

Next, we tested if the 5’ *CYSTM* α sequence enhances expression of a complex polycistronic expression transgene, named RUBY (28). RUBY consists of three betalain biosynthetic genes, *CYP76AD1, DODA* and a *Glucosyltransferase*, linked by two 2A peptides that are driven by a single promoter (28). We fused the 5’ *CYSTM* α sequence upstream of *CYP76AD1* coding sequence and transiently expressed the new transgene and the control vector in the leaves of *N. benthamiana*. At the 36- and 48-hour time points, we observed approximately a 1.5-fold increase in the leaf betalain content demonstrating the 5’ *CYSTM* α sequence enhances transgene expression from a complex polycistronic transgene (Fig. 1D).

### 5’ CYSTM α sequence enhances translation of *luciferase* in *A. thaliana* and TnT® Coupled Wheat Germ Extract System

Transient expression in *A. thaliana* leaves is often challenging due to the variable and low efficiency (29). Following a slightly modified protocol described by (30) we obtained consistent transient expression under our growth conditions. To quantify transgene expression, we used the Dual-Luciferase® Reporter Assay. We observed a 3-fold increase in firefly luciferase activity from both pluc 5’ *CYSTM* α and pluc 5’ *CYSTM1* constructs compared to the control, and a 2.2-fold increase compared to the pluc 5’ TMV Ω.

Additionally, in stable transgenic *Arabidopsis* plants we detected a 7-fold increase in the FLUC/RLUC ratio from pluc 5’ *CYSTM* α relative to the control, and a 2.3-fold increase compared to the pluc TMV Ω (Fig 2B). Notably, pluc 5’ CYSTM1 produced 60 % higher bioluminescence than the pluc TMV Ω but was 30 % lower the pluc *CYSTM* α. Since the enhancer effect of the full-length *CYSTM* 5’ UTR (pluc 5’ *CYSTM1*) was lower than that of the shorter 5’ *CYSTM* α sequence (pluc 5’ CYSTM α), we did not pursue it further.

**Figure 2:**
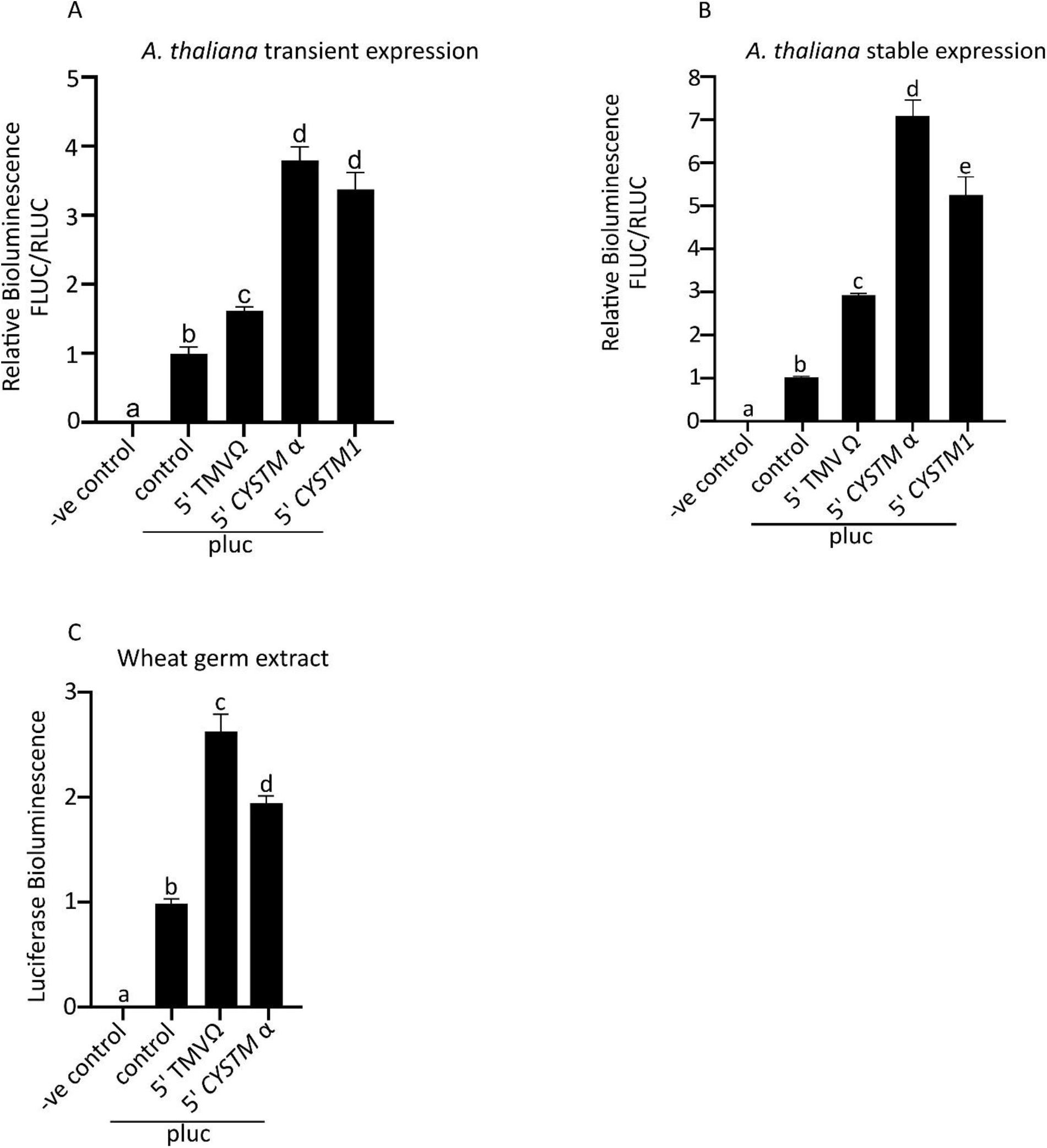
*CYSTM1* enhances dual luciferase expression in *A. thaliana* and TnT® Coupled Wheat Germ Extract System. **A)** Quantitative dual-luciferase assay results are shown for leaves expressing pluc 5’ *CYSTM* α compared to control constructs pluc and pluc 5’ TMV Ω. The experiment included five biological replicates and was repeated at least three times. Error bars represent the standard error of the mean. Statistical significance is denoted by letters (a–d, P < 0.01, one-way ANOVA). **B)** Quantitative Dual-Luciferase assay results from stably transformed T_1_ *A. thaliana* lines expressing different constructs. The experiment included five biological replicates. Error bars indicate the standard error of the mean. Statistical significance is indicated by letters (a– e) (P < 0.01, one-way ANOVA). **C)** Luciferase activity in an in vitro TnT® Coupled Wheat Germ Extract System for pluc (control), pluc 5’ TMV Ω, and pluc 5’ *CYSTM* α. Error bars represent the standard error of the mean. Statistical significance is indicated by letters (a–d) and was determined using one-way ANOVA (P < 0.01).

Previous studies have shown that enhancers effective in dicotyledonous plants often do not exhibit comparable efficacy in monocotyledonous species (3, 14). To evaluate the cross-species potential of our sequence, we performed the wheat germ assay. We observed a two-fold increase in LUCIFERASE protein from pluc 5’ *CYSTM* α compared to the control. In contrast, pluc 5’ TMV Ω exhibited only a 0.25-fold higher protein accumulation relative to pluc 5’ *CYSTM* α (Fig 2C).

### 5’ *CYSTM* α sequence enhances translation of firefly *luciferase* in *E. coli*

Previous studies have shown that enhancers of eukaryotic origin typically do not function effectively in prokaryotic systems due to fundamental differences in their translational and regulatory mechanisms (16–18). However, the TMV Ω sequence has been reported to enhance translation in both eukaryotic and prokaryotic organisms (19, 20). Building on these findings, we assessed the activity of pluc 5’ *CYSTM* α, which had previously demonstrated strong translational enhancement across plant species and reporter systems, including LUCIFERASE and RUBY, consistently outperforming TMV Ω. To evaluate its functionality in a prokaryotic context, we tested the construct in *E. coli* (Fig. 3A). Dual-Luciferase assay results revealed a 5-fold increase in relative bioluminescence for the pluc 5’ *CYSTM* α compared to the control, and a 38-fold increase for the pluc 5’ TMV Ω construct relative to the same control (Fig. 3B). These findings indicate that while the 5’ *CYSTM* α sequence enhances translation in *E. coli*, its activity is substantially lower than that of the TMV Ω.

**Figure 3:**
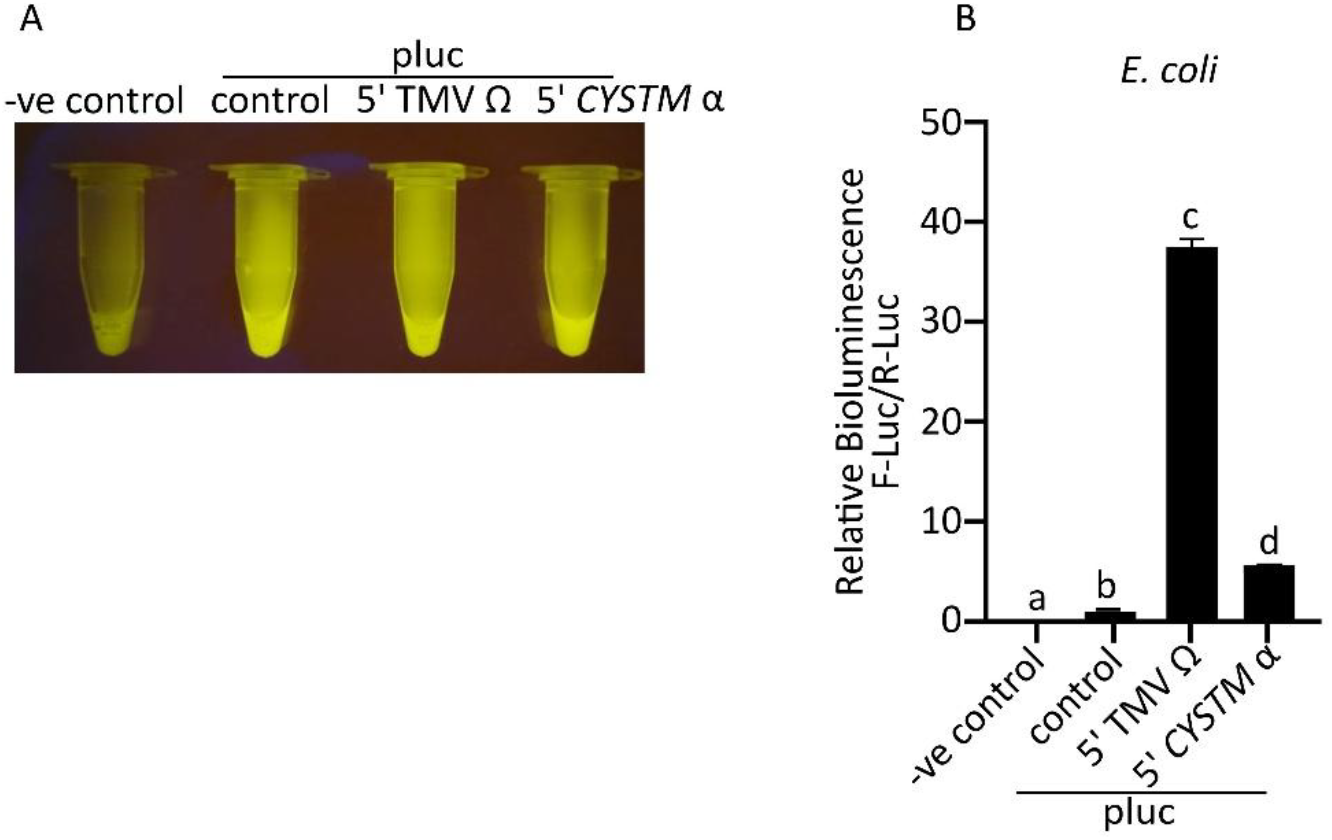
5’ *CYSTM* α enhances dual luciferase expression in *E. coli*. **A)** Qualitative luminescence imaging of *E. coli* transformed with pluc control, pluc 5’ TMV Ω and pluc 5’ *CYSTM* α. Colonies were grown to log phase, illuminated using a blueBox™ S Transilluminator and imaged with a Samsung Galaxy S20 FE. **B)** Quantitative Dual-Luciferase® Reporter Assay results showing relative bioluminescence from *E. coli* expressing pluc (control), pluc 5’ TMV Ω, and pluc 5’ *CYSTM* α constructs. Error bars represent the standard error of the mean. Statistical significance is indicated by letters (a–d) (P < 0.01, one-way ANOVA).

## Materials and methods

### Plant Materials

*A. thaliana* (Columbia ecotype) plants were cultivated in controlled environment rooms (Phoenix Biosystems), where the temperature was maintained at 21°C under Heliospectra ELIXIA LED lights with all 4 tuneable LED channels; 450 nm, 660 nm, 735 nm and 5700 K set to 200 µmol m^−2^ s^−1^. Plants were grown under long-day photoperiod conditions of 16 h light and 8 h darkness for 3 weeks. *N. benthamiana* (RDR6i) plants were germinated on soil in a master pot and then transferred to individual pots after 10 days. Plants were grown in Phoenix Biosystem controlled environment room with metal halide lights (100 µmol/m^2^/s) for 3 weeks.

### Plasmid Construction

Double stranded fragments containing either the full-length 110 nt 5’ UTR *CYSTM1* or a short 79-nt CYSTM α sequence flanked by SalI and PstI restriction sequences were synthesized as gBlocks by Integrated DNA Technologies (IDT), double-digested using SalI-HF (NEB) and PstI-HF (NEB) restriction enzymes, before cloning immediately 5’ of the *Firefly Luciferase* initiation codon of the dual luciferase plasmid pluc (pGrDL-SPb, Addgene #83205) (31). In p35S:RUBY (Addgene #160908), the 79-nt *CYSTM* α sequence was cloned immediately 5’ of the initiating *CYP76AD1* ORF (28).

### Transient expression by Agroinfiltration in *N. benthamiana* leaves

Transient infiltration was performed using a modified protocol of (32). Briefly, three mature leaves from 3-4-week-old *N. benthamiana* plants were selected for agroinfiltration. To ensure even distribution of the bacterial suspension (1X MS salt (4.3g/L); 10mM MES; 3% sucrose; 200uM acetosyringone, pH= 5.6), infiltration was performed over the entire leaf area via the abaxial surface using a 1 mL plastic syringe without a needle. Following infiltration, the plants were incubated in a controlled growth environment for two days prior to downstream experiments.

### Transient expression by agroinfiltration in *A. thaliana* leaves

*Arabidopsis* transient expression by agroinfiltration was performed according to a slightly modified protocol described by (30). The protocol was followed exactly up to the infiltration step however, an extended dark period of 60 hours instead of 24 hours was applied to the plants.

### Dual Luciferase assays

Quantitative dual luciferase assay was performed as per the protocol described by (32). Qualitative imaging of *N. benthamiana* leaves to detect luciferase activity was performed as described by (32) using Promega Dual-Luciferase® bioluminescence reporter system (Cat No: E1910).

### Dual Luciferase assay for *E. coli* cells

*E. coli* cells containing either the pluc control, 5’ TMV Ω or 5’ *CYSTM* α vectors were streaked onto LB plates from a −70 ^°^C glycerol stock, single colonies were transferred to LB broths and grown for approximately 12 hours at 37 ^°^C. The cultures were diluted to an OD_600_ of 0.2 and grown until the OD_600_ was 0.6. 1.5 mL of bacterial culture was pelleted at 14,000 x g for 3 mins, the supernatant discarded, and the pellet was resuspended in 50 µL of passive lysis solution provided in the Promega Dual-Luciferase® bioluminescence reporter system (Cat No: E1910) and 15 µL of this solution was used to perform the quantitative dual luciferase assays in a Promega GloMax® Discover according to the manufacturer’s protocol.

### Betalain (RUBY) extraction and quantification

Spectrophotometric Quantification of Betacyanins produced by RUBY in plant tissue was performed as per the protocol described by (33).

### RNA extraction, cDNA synthesis and Quantitative PCR

Total RNA was extracted from 5 mg discs of *N. benthamiana* leaf tissue using the Spectrum Plant total RNA kit (SIGMA-ALDRICH) and contaminating DNA removed using DNase (SIGMA-ALDRICH). cDNA synthesis for qRT-PCR was performed using oligo dT_20_ primer and an Invitrogen SuperScript IV kit as per the manufacturer’s instructions and PCR amplification was performed on an QuantStudio™ 7 Flex Real-Time PCR System (Applied Biosystems™). The oligonucleotides used in this study were F-Luc_F: 5’ TGCACATATCGAGGTGGACATC 3’, F-Luc_R: 5’ TGCCAACCGAACGGACAT 3’, R-Luc_F: 5’ CTGGCTCAATATGTGGCACAA 3’, R-Luc_R: 5’ CATGGTAACGCGGCCTCTT 3’. Amplification values were calculated using the ΔΔCT method (34).

### Protein analysis by western blotting

Proteins were separated on two SDS-PAGE gels, with one gel Coomassie-stained as a loading control and the other transferred to PVDF membranes (Mini Trans-Blot Cell, Bio-Rad) overnight at 4^°^C using standard transfer buffer (48 mM Tris, 39 mM glycine, 20 % methanol, 1.3 mM SDS, pH 9.2). Membranes were probed with anti-luciferase antibody (Sigma-Aldrich, L0159) and HRP-conjugated rabbit secondary antibody (Abcam, ab6721). Signals were detected with SuperSignal™ West Pico PLUS (Thermo Fisher, 34580) and imaged on a ChemiDoc XRS+ (Bio-Rad). Band intensities were quantified with ImageJ (NIH) and analysed using GraphPad Prism 9.

## Discussion

Maximizing transgene expression remains a central goal in plant molecular farming and synthetic biology (35). Translational control, particularly via 5′ untranslated regions (5′ UTRs), has emerged as a powerful strategy to enhance protein yields without modifying promoters or coding sequences (36). In this study, we identified a short, 79-nucleotide 5′ UTR fragment from the *A. thaliana CYSTM1* gene (5′ *CYSTM* α) that significantly enhances protein translation across multiple platforms, outperforming the widely used TMV Ω enhancer in *A. thaliana* and *N. benthamiana*, and exhibiting robust activity in *E. coli* and wheat germ extract.

The translational activity of 5′ *CYSTM* α performs better than other widely used plant and viral enhancers, including TMV Ω (20, 21), the alcohol dehydrogenase (ADH) 5′ UTRs from *A. thaliana, N. benthamiana*, and *O. sativa*, and synthetic constructs like synJ (22-24, 26, 37). In both transient and stable assays, 5′ *CYSTM* α conferred significantly higher expression than TMV Ω. Its effectiveness within a polycistronic RUBY construct (28) also supports its broader utility in multigene assemblies. These results position 5′ *CYSTM* α as a compact and potent regulatory element for enhancing protein expression in plant biotechnology.

Mechanistically, the enhancement mediated by 5′ *CYSTM* α appears to be post-transcriptional. Consistent with previous findings for TMV Ω and ADH-derived UTRs (20, 24, 26), qRT-PCR revealed no significant difference in transcript abundance, suggesting that translation efficiency is the primary driver of increased protein accumulation. TMV Ω’s activity is known to depend on poly (CAA) motifs that recruit the HSP101 chaperone and translation initiation factors (20, 38, 39). The 5′ *CYSTM* α sequence similarly contains 3 (CAA) repeats, 1 AMAYAA motif and 2 putative m^6^A methylation motifs (DRACH), which have been implicated in enhanced translation initiation via eIF3 binding in other systems (40-45). Interestingly in publicly available m^6^A-SAC-seq data, one of the two m^6^A motifs was methylated implicating enhanced translation initiation via eIF3 binding (46). Furthermore, using publicly available ribosome profiling data of *Arabidopsis* (47) revealed that *CYSTM1* had 1132 ribosome footprint reads across the 216-codon ORF, corresponding to 758 P-sites, of which approximately 90 % were in-frame. Comparison with RNA-seq coverage indicated a translation efficiency of approximately 3.25, consistent with highly efficient translation. The high density of ribosome footprints along the coding sequence and the predominant in-frame P-site distribution collectively suggest that the entire ORF is robustly and actively translated. These features may contribute to its functionality and warrant further investigation through structure-function dissection, RNA-binding assays, and m^6^A mapping.

A particularly intriguing aspect of 5′ *CYSTM* α is its cross-kingdom activity. While plant-derived enhancers typically fail in prokaryotes due to fundamental differences in translation initiation (48-50), 5′ *CYSTM* α increased luciferase expression five-fold in *E. coli*, though it was less effective than TMV Ω. This residual activity suggests an overlap with bacterial translation signals, possibly through stable RNA structures or Shine-Dalgarno-like motifs (51, 52). This ability to work in both plant and bacterial systems shows that the 5′ *CYSTM α* sequence could be a flexible tool for building synthetic biology systems in either plant or microbial hosts.

Practically, 5′ *CYSTM* α complements existing strategies to enhance gene expression, such as promoter engineering (53), intron-mediated enhancement (12, 54) and codon optimisation (13). Its small size and high efficacy make it particularly suitable for synthetic circuits and compact vectors. Applications may include boosting yields in plant-based biomanufacturing (55), enhancing biosynthetic pathway flux (56), or increasing sensitivity of reporters in functional genomics studies (23).

Future work should aim to identify the minimal active region, assess structure–function relationships, and test for the presence of m^6^A modifications and protein binding. Ribosome profiling and cross-species testing will further refine our understanding of its mechanism and utility. In summary, 5′ *CYSTM* α is a post-transcriptional enhancer that improves protein yield across diverse hosts. Its cross-kingdom functionality and compatibility with multigene constructs underscore its potential for use in plant and microbial synthetic biology.

## Data Availability

All plasmids generated in this study have been deposited in Addgene under the following accession numbers: 247490, 247491, 247492, 247493, and 247568.

## Acknowledgements

This research was supported by an Australian Research Council (DP190101303) awarded to I.R.S. and a Research Training Postgraduate Scholarship was awarded to J.K. The authors also acknowledge the use of ChatGPT (GPT-5) and GPT-4-Turbo for suggestions in improving writing style, outline structure, proofreading of the manuscript and providing high-level feedback. No content generated by AI technologies has been presented as our own.

## Conflicts of interest

The authors declare no conflict of interest.

## Supplementary Data

**Supplementary figure 1:**
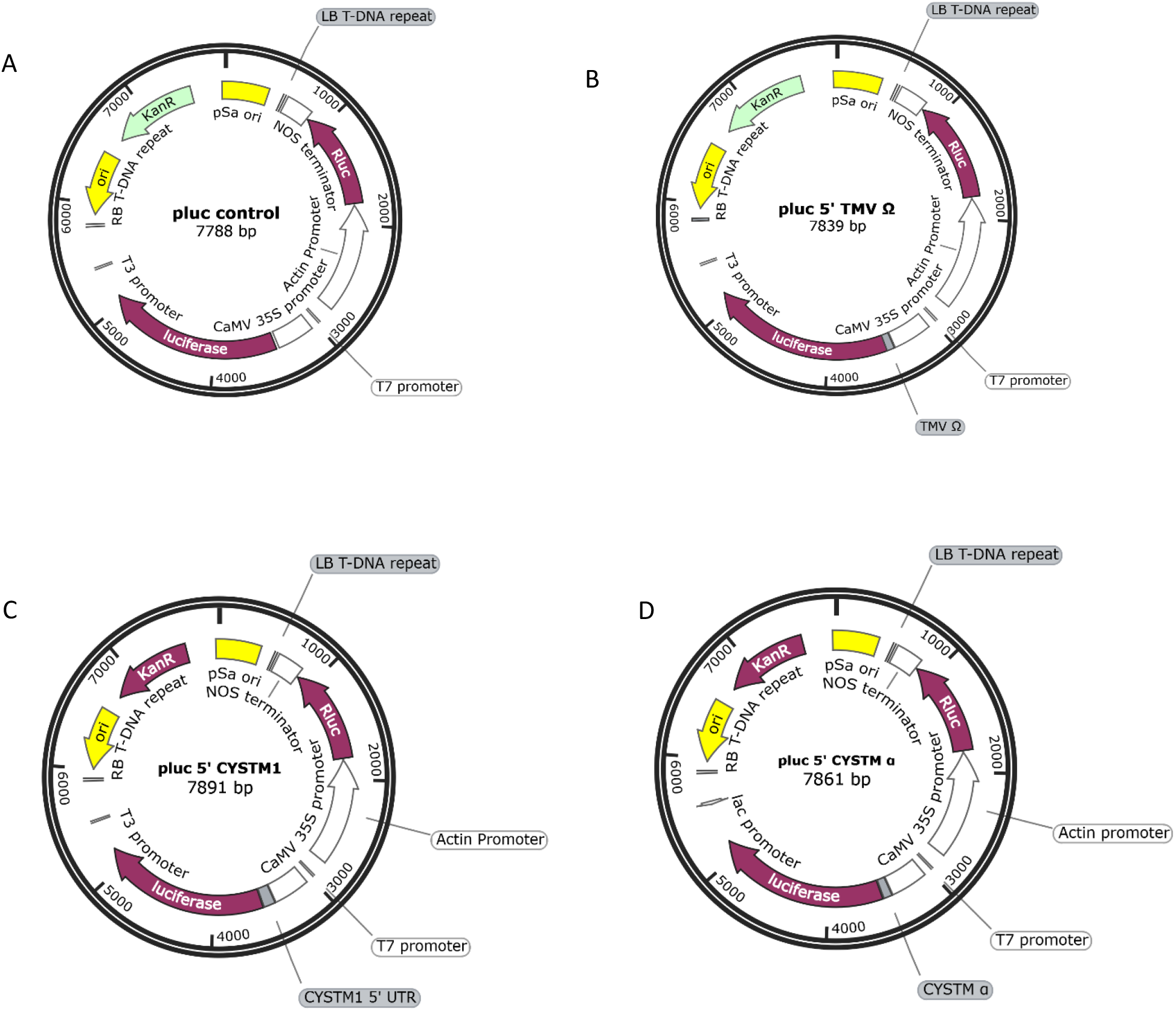
Plasmid maps of, **A**) pluc control, **B**) pluc 5’ TMV Ω, **C**) pluc 5’ CYSTM1 and **D**) pluc 5’ CYSTM α.

**Supplementary figure 2:**
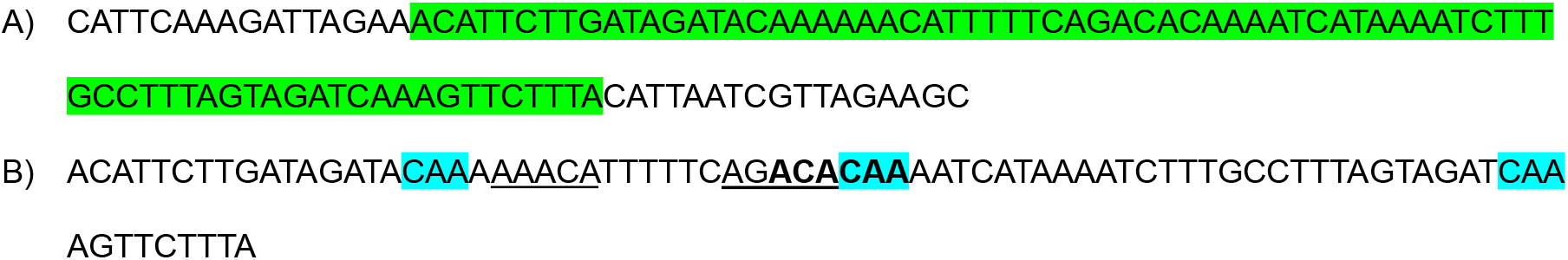
**A**) Complete 5’ UTR of the *CYSTM1* gene with 5’ CYSTM α sequence highlighted in green. **B**) 5’ CYSTM α sequence with CAA motifs highlighted in blue, DRACH motifs underlined and AMAYAA motif in bold.

## Notes

### Competing Interest Statement

The authors have declared no competing interest.

### Summary of Updates

Figures 1 and 3 have been updated to remove artifacts generated due to low resolution PNG images.

